# Photon-Resolved Excitation-Denoised (PRED) Three-Photon Imaging Improves Detection of Neuronal Activity in Awake and Behaving Mice

**DOI:** 10.64898/2026.04.10.717694

**Authors:** Tiberiu S. Mihaila, Eunji Kong, Adrian Negrean, Tristan Geiller, Darcy S. Peterka, Attila Losonczy

**Affiliations:** Mortimer B. Zuckerman Mind Brain Behavior Institute, Columbia University, New York, NY, United States; Peter O’Donnell Jr. Brain Institute, University of Texas Southwestern Medical Center, Dallas, TX, United States; Department of Neuroscience and Wu Tsai Institute, Yale University, New Haven, CT, United States; The Kavli Institute for Brain Science, Columbia University, New York, NY, United States

## Abstract

Three-photon microscopy (3PM) has enabled the optical access of neurons ∼500-1500 µm below the brain surface but has been limited to slow imaging frame rates or small total imaged areas by competing constraints: a high peak power requirement for nonlinear excitation, and the need to limit total average power. Additionally, the high sensitivity to laser power fluctuations and the inherently dim signals introduce additional challenges and add error. When combined with other effects, like brain motion in behaving animals, 3P imaging of neuronal activity during animal behavior has remained inadequate. Herein, we systematically address these limitations by 1) carefully balancing scanning speed with power requirements, 2) using a deeply cooled silicon photomultiplier detector and Bayesian statistics-based processing (PRED-correction) to reduce excess noise, and 3) by spatiotemporal shaping of excitation pulses. Our improvements enable rapid (20-30Hz) imaging of calcium activity in the dorsal hippocampal dentate gyrus of behaving mice, allowing the identification of spatially tuned neurons and the recapitulation of established functional properties across different cell types in this brain region. PRED-3P imaging provides a new approach to functional characterization of cells deep in the brain that were previously inaccessible to two-photon imaging.

## Introduction

Multiphoton fluorescence imaging is invaluable for understanding neuronal structure and function in the brains of behaving animals. The advent of two-photon microscopy^1,2^ (2PM) has enabled *in vivo* optical recordings hundreds of micrometers deep within scattering brain tissue. Two-photon (2P) excitation requires near-simultaneous absorption of two photons,^3,4^ with a quadratic dependence on light intensity. When used in imaging, this confers the highest excitation efficiency near the objective’s focal point and limits both photodamage and bleaching outside of the area of interest. This combination has enabled countless cellular and subcellular measurements of neuronal activity across the brain during diverse animal behaviors. However, 2PM is generally limited to regions ≤∼600µm from the surface of the brain^5^ because scattering reduces the peak power at the focus exponentially and when measuring with the high power needed for imaging at depth, the power at the surface generates background signal comparable or higher than that at the focus. As depth increases, scattering and brain-surface background accumulate to reduce contrast below a usable level.^6^ Three-photon microscopy (3PM) overcomes this by using longer excitation wavelengths that scatter less in biological tissue and which are lower in energy, requiring simultaneous absorption of three photons to excite fluorescent molecules.^7^ The increased non-linearity improves contrast and yields roughly doubled depth-of-penetration,^8^ enabling recordings at ∼1300µm depth, bringing 2P-inaccessible cells into view^9-14^

Nevertheless, applying 3PM to biology faces its own challenges in 1) excitation light delivery and 2) in emitted photon detection. The cubic nonlinearity demands high peak light intensities, while the living tissue requires that average power is below the threshold where thermal damage occurs. This has meant that short (≤75-fsec) excitation pulses are used at relatively low repetition rates, maximizing power per pulse. Each pixel requires at least one laser pulse to generate signal, and for a given deposited power per pixel, single pulses generate signal more efficiently than splitting that power across multiple pulses.^15^

Most 3P experiments have used lasers tuned to ∼0.5-2 MHz repetition rates; however, for large field of view (FOV) or high-resolution imaging, this pulse number limit restricts the total pixel rate, leading to imaging frame rates below those needed to benefit from modern fluorescent calcium sensors. This trade-off has severely limited 3PM in functional imaging, with only a single study imaging a FOV wider than 250 µm at 10 Hz.^16^ Furthermore, the single-pulse-per-pixel strategy inevitably complicates measurements as the shot-to-shot fluctuations of the laser pulse intensities^17^ are amplified through the cubic non-linearity of the excitation process. Finally, with all deep imaging regardless of excitation strategy, the magnitude of changes in dimmer signals from neural activity fall near the scale of the Poisson variation that is intrinsic to any point emitter. Any excess noise from the system will further hamper the ability to detect changes in fluorescence.

Here, we report a 3PM implementation that solves these problems, enabling high-frame rate, wide-FOV fluorescence population calcium imaging of hundreds of neurons >600µm below the brain surface in behaving animals. We show that 3P-excitation at ∼4 MHz can be coupled with rapid resonant-galvo-galvo scanning to perform functional imaging over an area of ∼250 µm × 350 µm at ≥20Hz. We image the activity of deep neurons in the dentate gyrus (DG) of the dorsal mouse hippocampus through an intact overlying hippocampal area CA1, capturing granule cells of the lower blade of this structure for the first time *in vivo*. By using a deeply cooled silicon photomultiplier detector coupled to a new Bayesian-statistics-based photon-counting approach, we eliminate most excess noise, and by optimizing the spatiotemporal profile of the excitation pulse, we maximize the number of detected photons per pulse. With these improvements, we recapitulate known properties of different DG cell types during animal behavior, identifying spatially tuned mossy cells, without damaging the overlying CA1 region and compromising information output from the hippocampus.

## Results

### Fast laser-repetition rate resonant-galvo-galvo 3P imaging enables large-area, high-speed imaging

To better understand the trade-off between imaging power and frame rate, we modeled the combinations of laser-repetition and scanner line-rates that enable high frame rates in different FOV sizes **(Fig. 1A)**. To image a FOV of a given area at a desired frame rate, the scanner must achieve a minimally required speed. Most reported 3PM implementations have been scan-limited systems, using galvanometer-galvanometer (GG) scanning modules, with typical line rates of <1kHz, constraining large-FOV 3PM frame rates to 5-10Hz. As the scanner speed is increased, a threshold is passed above which the imaging speed instead becomes limited by the repetition rate of the laser since each pixel must receive at least one excitation pulse, setting the maximum number of pixels per line. At higher repetition rates, each laser pulse contains less energy at a given average imaging power. However, 3P-excitation efficiency depends strongly on peak pulse energy, typically requiring >1 nJ at the focus for successful imaging (∼15-50kW at the surface)^18-19^ (**Fig. 1B**). This peak power floor mandates an increase in average laser power with repetition rate. This requirement increases with depth, as non-zero tissue scattering and absorption attenuate the pulse power with depth. If the average laser power is raised too much, the risk of thermal damage increases.

**Figure 1.**
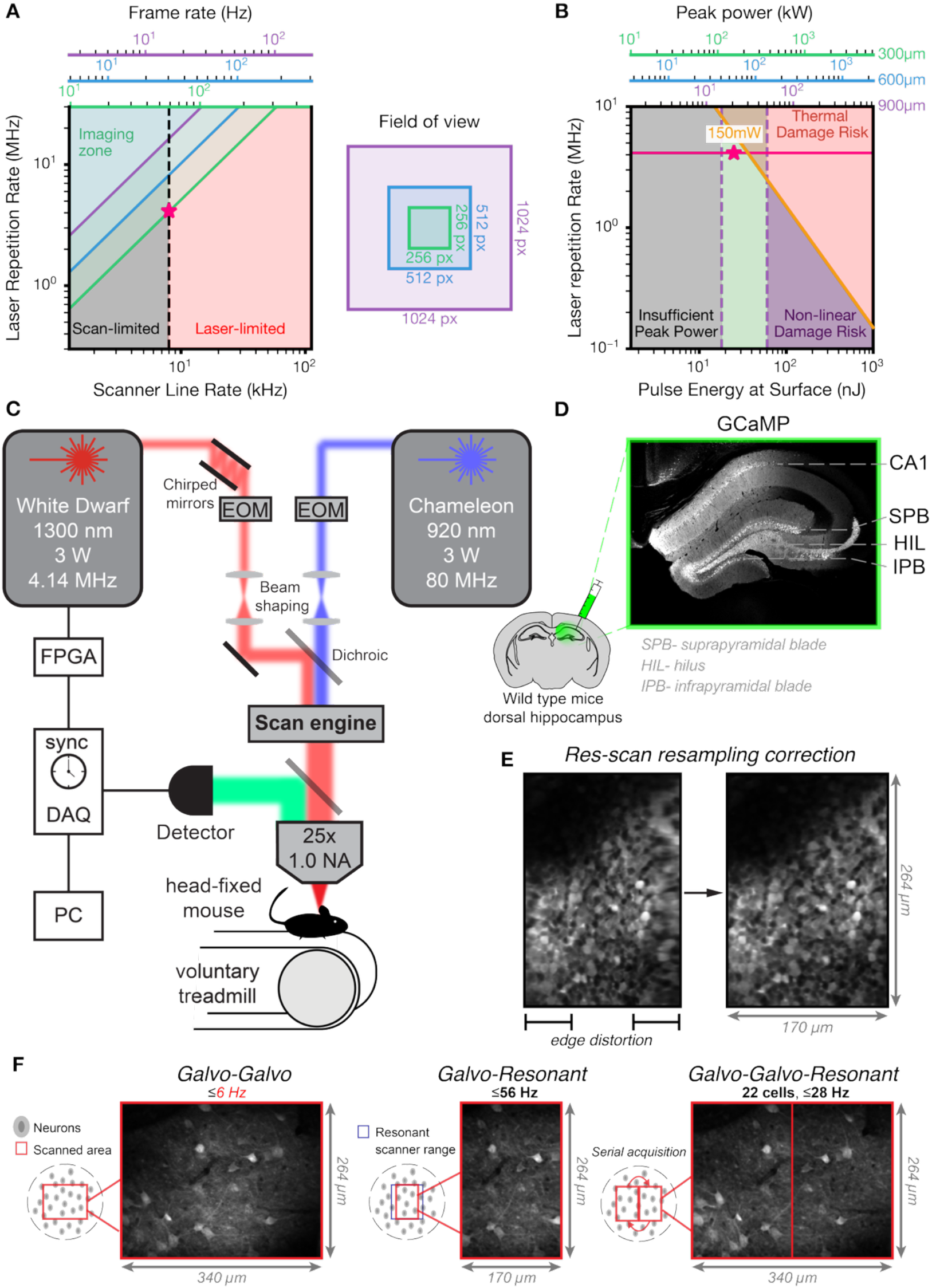
Rapid scanning and high laser repetition rates are required to achieve >20-Hz 3P imaging. **A)** Calculated laser repetition rates and scanner line rates required to achieve imaging at various frame rates in different-sized regions of interest (ROIs). Pink star indicates parameters utilized in our system. **B)** Calculated laser repetition rate regime in which sufficient peak power for 3P imaging can be achieved at various depths, without crossing the thermal damage threshold. Pink star indicates parameters utilized in our system. **C)** Overview of integrated multiphoton microscope built in this study. **D)** Example calcium indicator fluorescence following adeno-associated viral infection of the dorsal mouse hippocampus. **E)** Resonant scanner resampling correction eliminates spatial distortions in the acquired images. *Field-of-view* (*FOV) size – 170 µm × 264µm* **F)** Resonant-galvo-galvo scanning can image a given FOV at much faster frame rates than galvo-galvo scanning and can image a wider area than galvo-resonant scanning.

We identified a narrow window between these constraints, wherein the use of a ∼4MHz laser repetition rate with an 8kHz scanner would allow 20-30Hz imaging of a ∼(300-500µm)^2^ FOV, while limiting both the peak energy and the total thermal load below their damage thresholds (**Fig. 1A, B**). To test the effective performance *in vivo*, we constructed a combined multiphoton imaging system featuring a 1300-nm 3P excitation laser with a 4.14MHz pulse repetition-rate, as well as a conventional tunable 2P excitation laser at 80MHz. The excitation beams are co-aligned into the microscope and steered by a resonant galvo-galvo scan engine, which can perform both traditional 1kHz galvanometer line scanning and rapid 8kHz resonant line scanning (**Fig. 1C**).

We applied our system in the DG of the dorsal murine hippocampus. With cells ∼600-1000µm below the top of the dorsal hippocampal CA1, DG has remained the least accessible hippocampal area for 2PM. Published studies have mostly imaged this region by removing CA1, compromising the output of hippocampus. They have primarily focused on the suprapyramidal blade (SPB) of DG,^20-26^ with a paucity of work reported in the DG hilar region (HIL), ^27-28^ and none in the infrapyramidal blade (IPB) which contains half of the of DG cell population.^29^ To label cells, an adeno-associated virus expressing the genetically encoded calcium indicator jGCaMP8s^30^ was stereotactically injected throughout the entire depth of the dorsal hippocampus, enabling expression of the fluorescent indicator from the deepest region (IPB) to the superficial CA1 pyramidal neuron layer. (**Fig. 1D**). A chronic imaging window was then implanted above CA1, and mice were imaged after ≥2 weeks while head-fixed on a treadmill under the objective lens of our microscope.

To maximize the FOV, each pixel received a single laser pulse and was sampled phase-locked with the laser reference clock and integrated for a ∼12.5-nsec window after each pulse. Given that the resonance scanner velocity is sinusoidal, but the laser clock is constant in time, the center of the scan field is less-densely sampled in pixel space (and thus spatially compressed), while the edges are more frequently sampled (and thus stretched) (**Fig. 1E, left**). To correct this, we developed a resampling algorithm that reconstructs the image in physical space by fitting phase offsets and by using the positions of the pixel centers and the laser pulses to effectively remove sampling distortion across the FOV (**Fig. 1E-right, supplemental discussion 1**). Rapid resonant-GG scanning achieved sufficient imaging contrast at much faster imaging rates than galvo-galvo scanning (56Hz vs. 6Hz) but was limited to a narrower FOV (**Fig. 1F**), given that our combination 4MHz laser repetition rate + 8kHz scanner system is laser-limited at these imaging parameters (**Fig. 1A**). However, with the high line-rate of our system, we used the second galvanometer in our scan-head to move the FOV to an equal-width frame immediately adjacent to the FOV imaged by the galvo-resonant scanner (**Fig. 1F**). The strategy enabled in vivo 3P imaging at a 28 Hz framerate in a wide FOV.

### Rapid 3P imaging detects more cells than 2P imaging throughout the mouse dentate gyrus

We next compared the capabilities of 28Hz 3PM with what could be achieved by 2PM in a wide FOV at various depths through intact CA1. Thus, we recorded jGCaMP8s activity in four different regions of the dorsal mouse hippocampus: CA1, SPB, HIL, and IPB. We first acquired a *z*-stack through the dorsal mouse hippocampus, beginning superficially in CA1 and extending down approximately 900µm to the IPB blade of the DG. Indeed, 3P excitation achieved higher contrast at depth in the DG than 2P excitation, enabling clear visualization of many more cells (**Fig. 2A**). To quantify these improvements, we imaged at least two FOVs from each hippocampal sub-region for two-minutes in three head-fixed mice on a clamped burlap belt (**Fig. 2B**), first with 2P and then with 3P at matched average powers. We asked how many cells could be identified in an unbiased fashion from the different subregions. Cells were automatically segmented with the stock model (untrained) of CellPose from the mean image of the motion-corrected image time series. As expected, in the superficial CA1 region there was no significant difference in the number of identified pyramidal neurons (PN) or in the mean-image contrast with 2PM and 3PM (**Fig. 2C+D**). However, at depth in the DG, 3PM enabled identification of more granule cells in the SPB and IPB than 2PM and increased the contrast in every subregion of the structure (**Fig. 2C+D**). Furthermore, calcium transients were detected in 3PM recordings across all hippocampal regions, including the IPB, for which there are no reports of multiphoton imaging to date (**Fig. 2B iv**).

**Figure 2.**
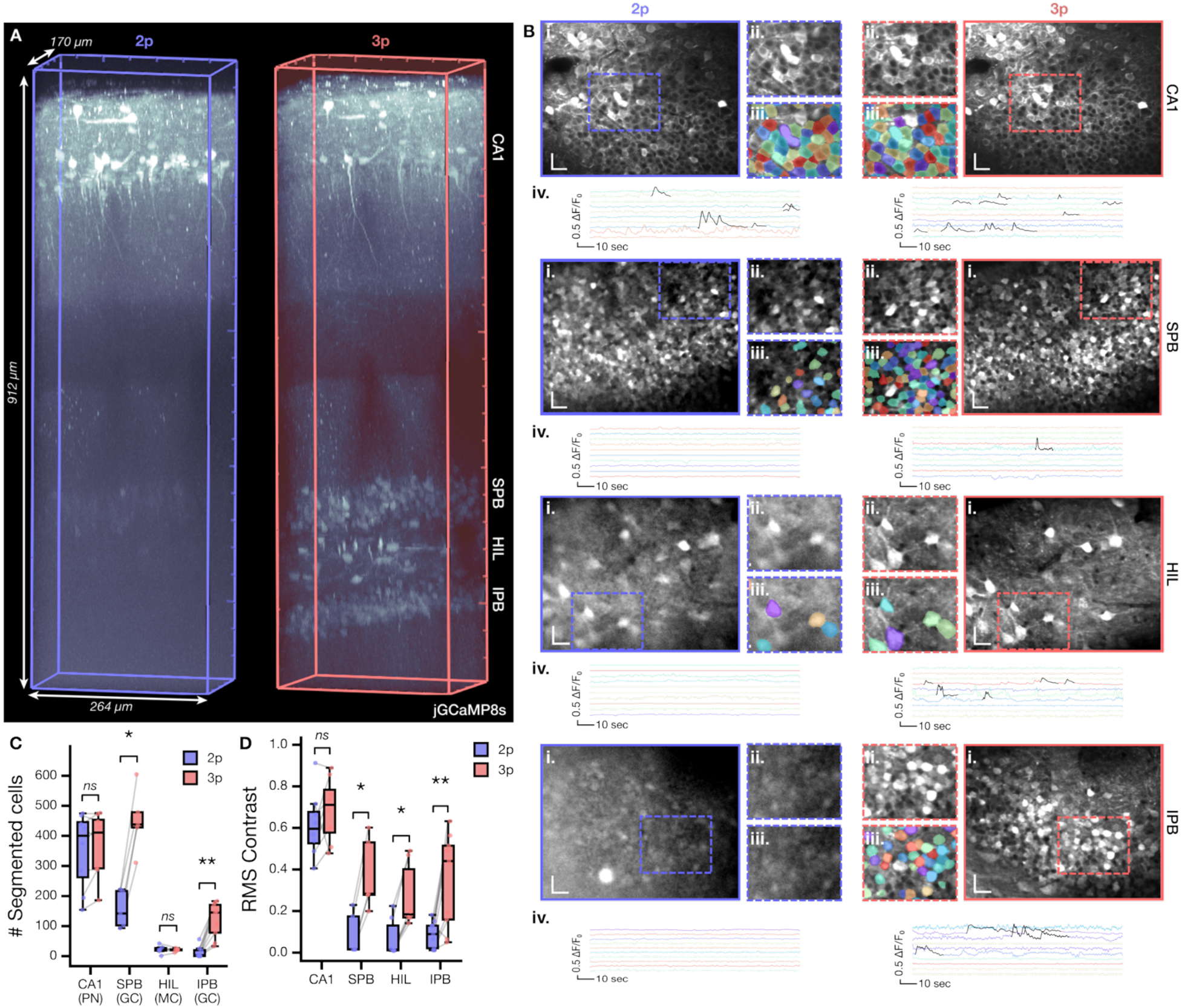
3P excitation enables the detection of more cells in the dorsal mouse hippocampus than 2P excitation. **A)** *In vivo z*-stack through the dorsal hippocampus of an anesthetized mouse infected with rAAV-jGCaMP8s. *Scale – 170µm* x *264µm* x *912µm* **B)** *i*. Mean images from serially acquired two-minute 2P & 3P recordings of the same FOV in hippocampal CA1, suprapyramidal blade (SPB; 607-652µm deep), hilus (HIL; 668-701µm deep), and infrapyramidal blade (IPB; 726-822µm deep). *ii*. Magnified region from the recorded FOV indicated in *i. iii*. Automatically drawn ROIs identified by untrained stock CellPose model. *iv*. Example calcium traces from recorded cells in each region with each imaging modality. *Scale-bars = 25µm*. **C)** Comparison of CellPose-detected cells in each region with 2PM vs. 3PM. Significance via Wilcoxon test. *p<0.05, **p<0.01. **D)** Comparison of root-mean square (RMS) contrast of the mean over the recordings with 2PM vs. 3PM. Significance via Wilcoxon test. *p<0.05, **p<0.01.

### Photon resolution and excitation decoupling (PRED) correction minimizes measurement error

Functional imaging measurements, e.g., identification of calcium transients, are complicated in the low photon flux regimes that are typical of efficient 3P imaging, because the magnitude of the intrinsic Poisson noise with photon detection can be on similar magnitudes to calcium-driven fluorescence changes. Thus, any extra measurement error or noise, which can originate from several sources, must be eliminated.

Additional error can arise from signal detection, as electronics invariably introduce noise. The short gated 12,5-nsec post-pulse collection window partially mitigates this, as most GCaMP photons should arrive in this window, given its fluorescence lifetime (**Fig. 3B**), and total dark counts are restricted compared to ungated integration. Nevertheless, electronic noise during the collection window cannot be controlled. Perhaps even more fundamental, given that only one pulse is delivered to each pixel, any shot-to-shot variation in laser pulse energy will affect the generated fluorescence. 3P excitation efficiency goes with the cube of the pulse energy (assuming no temporal envelope changes), so, e.g., 5% upward fluctuation results in a 16% increase in brightness **(Fig. 3B)**.

**Figure 3.**
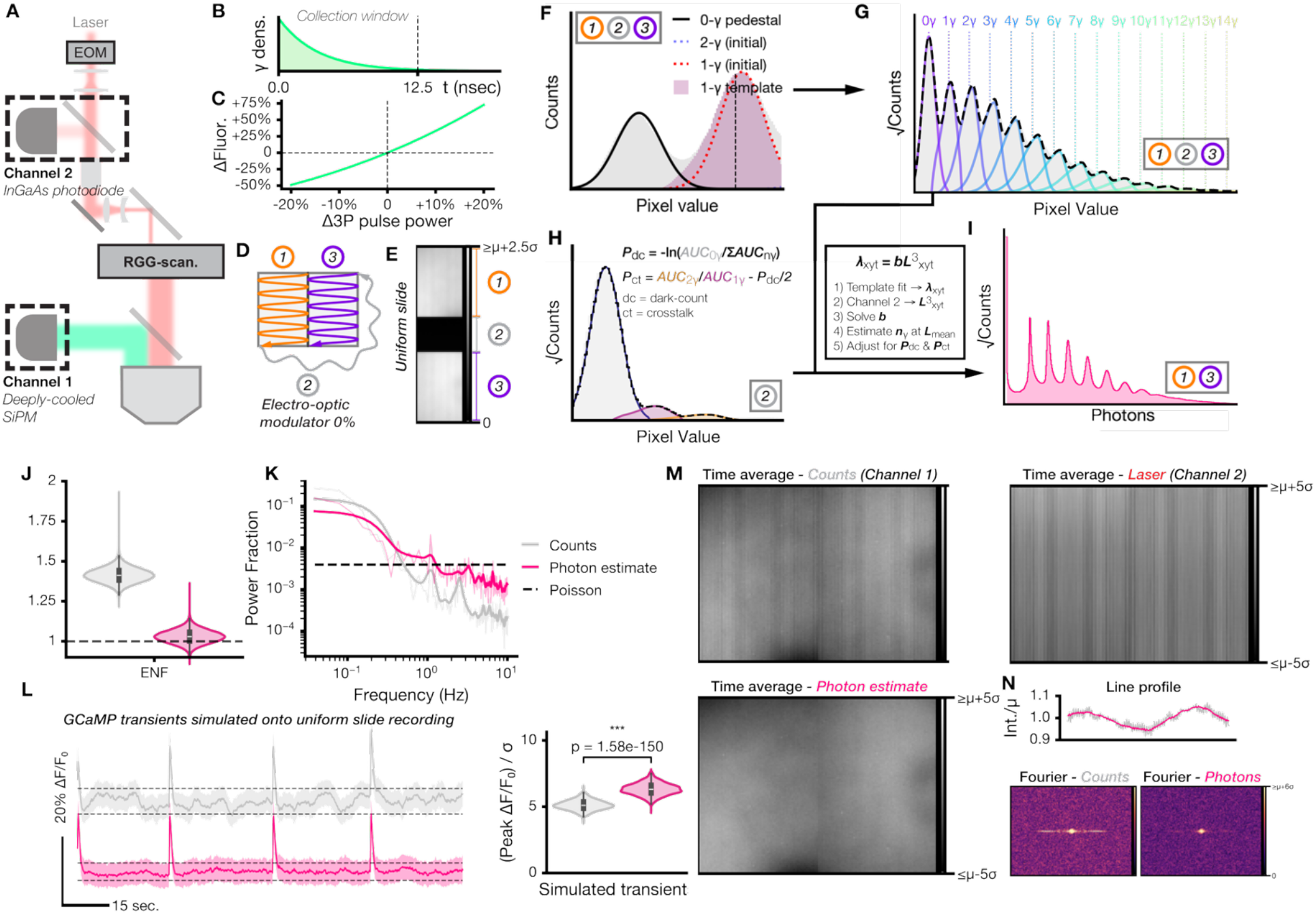
Photon resolution with excitation decoupling (PRED)-correction reduces excess non-Poisson noise in 3P-recordings. **A)** Detection strategy for emitted photon counting and laser pulse power measurement. **B)** Fluorescence was collected during a 12,5-nsec window synchronized to the laser pulse, during which most emitted GCaMP photons will arrive. **C)** Relationship between expected change in fluorescence vs. 3P-excitation fluctuation magnitude. **D)** Overview of measured regions in resonant-galvo-galvo 3P imaging. **E)** Example recorded data from the measured regions from a recording of a uniform fluorescent slide. **F)** 0- and 1-photon templates can be fit to the pixel histogram to model the detector response profiles to these events. **G)** Convolution of the 1-photon template generates templates for multi-photon detections that successfully fit the measured pixel histogram. **H)** 0-, 1-, and 2-photon templates can be used to estimate detector dark counts and crosstalk from the recorded dark region between ROIs. **I)** Photon template fitting and estimated dark-count & crosstalk probabilities can be used to estimate photon counts per pixel using a Bayesian inference model and photon statistics and to correct for pulse-to-pulse variation in laser power. **J)** Photon-estimate recording via PRED correction reduces the expected noise factor close to the Poisson limit as compared to gain-matched digitizer counts. **K)** Frequency response of photon-estimate recording more closely matches that expected of a Poisson distribution than that of the measured digitizer counts. **L)** PRED-correction reduces systemic fluctuations in the fluorescent baseline and enables significantly higher signal-to-noise ratio in simulated GCaMP transients. Averaged traces across 100-pixel ROIs (error 95% confidence interval). **M)** PRED-correction eliminates spatial fluctuations in excitation laser intensity. μ= mean pixel value. σ= standard deviation of pixel values. **N)** Intensity fluctuations in the scanning direction from electro-optical modulator ringing are eliminated by the PRED correction. *Top –* line profile in the scan direction. *Bottom –* Fourier transform of uncorrected counts vs. corrected photons. μ= mean pixel value. σ= standard deviation of pixel values.

To reduce noise from electronics and laser fluctuations, we made two modifications: 1) we replaced the photomultiplier tube detector with a deeply-cooled large-area silicon photomultiplier (SiPM) array — a detector type that scales highly-linearly with the number of incident photons, and 2) we measured and pixel-matched the instantaneous power of each laser pulse by sampling the beam with a beam pick-off and directly measuring the individual pulse power with an InGaAs photodiode (**Fig. 3A, S1**). Historically, large-area SiPMs have been limited by the rate of dark counts (detection events in the absence of incoming photons) and by pixel crosstalk (the probability that a detection event triggers an offspring pseudo-event). Recent advances have improved both these parameters, so we tested whether we could use the expected photon-scaling from these detectors to correct for electronic noise and laser power fluctuations in our recordings.

We performed resonant-GG imaging of two ROIs from a uniform slide (**Fig. 3D,E**), with our primary imaging channel recording the signal from the SiPM and the secondary channel being the instantaneous laser power as measured by our photodiode. Our scanning paradigm provides pixel information from three regions: 1) the first ROI, 2) the inter-ROI region passed by the beam with power floored via the electro-optic modulator, and 3) the second ROI (**Fig. 3D,E**). Given that there is nearly a million-fold reduction in 3P excitation efficiency in region 2 (typical contrast ratio from our Pockels Cell is ∼90:1), this provides a within-recording way to estimate the dark-count and crosstalk probabilities. Firstly, to characterize the response distribution from single-photon events, a Gaussian is fit to the zero-photon pedestal and to the peaks corresponding to one-photon (1-γ) and 2-γ events (**Fig. 3F**). Then, a 1-γ response template is fit, maximizing the goodness-of-fit to the pixel histogram when added to the 0-γ pedestal (**supplemental discussion 2.1**). Given the nonzero SiPM output pulse duration, photons that arrive later in the collection window trigger fewer digitizer counts within the 12,5-nsec collection window, causing the 1-γ response to approximate a leftward exponentially modified Gaussian rather than a simple normal distribution. Next, a 2-γ response template is generated by convolving the 1-γ template with itself, and additional multi-photon templates were generated by further convolution. These templates successfully fit the measured pixel distribution (**Fig. 3G**). The dark count and crosstalk probabilities are then calculated using the 0, 1-, and 2-γ templates in the recorded dark inter-ROI region (**Fig. 3H, supplemental discussion 2.2**).

With the dark-count + crosstalk probabilities and the photon-detection response templates in hand, we used Bayesian statistics to infer the number of photons measured in each pixel and to correct for bias introduced by fluctuations in the excitation laser (**supplemental discussion 2.3-2.6**). Briefly, we modeled the photon flux in each pixel as a function of its intrinsic brightness and the cube of the laser power, accounting for the probability of dark counts and crosstalk (**Fig. 3D-F**). We then inferred the photon flux at each pixel in each frame using the photon-response templates and the laser power at that pixel, as measured by the photodiode in channel 2. By modeling the pixel histogram at each point across the recording as the sum of Poissonian photon-flux distributions scaling with the cube of the laser power, the most likely baseline brightness of each pixel was solved. With the baseline brightness, we then used Bayesian statistics to estimate the expected detected photon number, and then the expected true fluorophore brightness, i.e. the photon number had the laser been perfectly stable at the recording average. By applying this approach to every pixel, we obtain a more discrete pixel distribution, approximating a photon-counting process (**Fig. 3I**).

This photon resolution and excitation decoupling (PRED) correction dramatically reduces the expected noise factor (ENF) compared to the original recording (gain-matched), bringing it very near the ENF of 1 expected for a Poisson process (**Fig. 3J**). Furthermore, the frequency response of the recording flattens, becoming more like that if Poisson/emitter noise were the only error source present (**Fig. 3I**). This correction also removes intensity patterns caused by spatial variation in the laser excitation power due to ringing of the electro-optical modulator (**Fig. 3M**). Fluctuations in the scan direction are nearly eliminated (**Fig. 3N**). Given that our correction is being applied on a per-pixel basis in time, treating each pixel as an independent emitter, the elimination of systematic average spatial differences between pixels demonstrates the robustness of the PRED correction.

To understand whether the PRED correction would improve calcium-indicator transient detection, we injected simulated GCaMP transients into our recording every 30-seconds. We then divided the uniform slide into equally sized 100-pixel ROIs and compared the distribution of traces across the ROIs. Slow and large-scale oscillations in the baseline that correlated with the laser were largely eliminated by the correction (**Fig. 3L, left**). This led to a significant increase in the transient amplitudes (**Fig. 3L, right**) and in detectability (**Fig. S2A**), enhancing the theoretical ability to identify calcium events. Indeed, comparison of PRED-corrected and non-corrected jGCaMP7f^32^ recordings during animal behavior provides further evidence that PRED enhances calcium transient detection (**Fig. S2B**,**C**).

### Optimization of spatiotemporal beam-shaping

Having optimized the detection side of our microscope with the PRED-correction, we next aimed to augment excitation efficiency by optimizing the spatial and temporal shapes of the pulses. To achieve the most spatially tight focus possible, a high-NA objective should be used with maximal backfilling (backfilling proportion, *β*=1). However, for *in vivo* multiphoton imaging, underfilling is common, as the total power coupling from laser to sample increases, the reduction of signal with depth from scattering and aberrations is reduced, and the slightly increased axial extent of the point-spread function partially mitigates the effects of *z*-motion during recording sessions.^2^ However, it has been argued that even narrower *β*-values (0.6-0.7) must be used for 3P imaging, as ∼1300-nm light is absorbed by water in tissue significantly more than light at wavelengths typically used for 2P excitation, and therefore longer optical path lengths from a fully backfilled objective become much more attenuated than the narrower angle light (which travels a shorter distance), reducing the effective illumination efficiency. In fact, simulations suggest that the deeper one images, the lower the optimal *β* drops.^5,31^ Nevertheless, this has never been tested in a real biological sample.

To measure the effect of objective backfilling on 3P excitation efficiency in the hippocampus, we implemented a variable magnification telescope (**Fig. 4A**) that enabled us to achieve backfill-proportions of 0.6, 0.7, and 0.9 (**Fig. 4B**). We took 30-second average-power matched recordings from the SPB, HIL, and IPB in the dorsal DG of three mice with different *β* and quantified the change in detected photons per pixel versus the average of all recordings as a function of *β* (**Fig. 4C**). Surprisingly, we found no difference in any of the regions, suggesting that similar focal powers were achieved, regardless of *β*. Given that backfilling also affects resolution, we aimed to determine whether *β* affected automated cell detection in these hippocampal regions. As expected, the most highly compressed beam (*β*=0.6) resulted in significantly fewer cells in the deeper regions, consistent with the inverse relationship between backfilling and resolution. Furthermore, we found that the most expanded beam (*β*=0.9) also dropped off in the deepest region, the IPB, perhaps due to slightly higher scattering of the longer lateral light paths to the focus. Thus, in our hands, the optimal backfilling ratio, irrespective of imaging depth, was 0.7, which provided the best resolution and lowest falloff (**Fig. 4D**) and was used moving forward.

**Figure 4.**
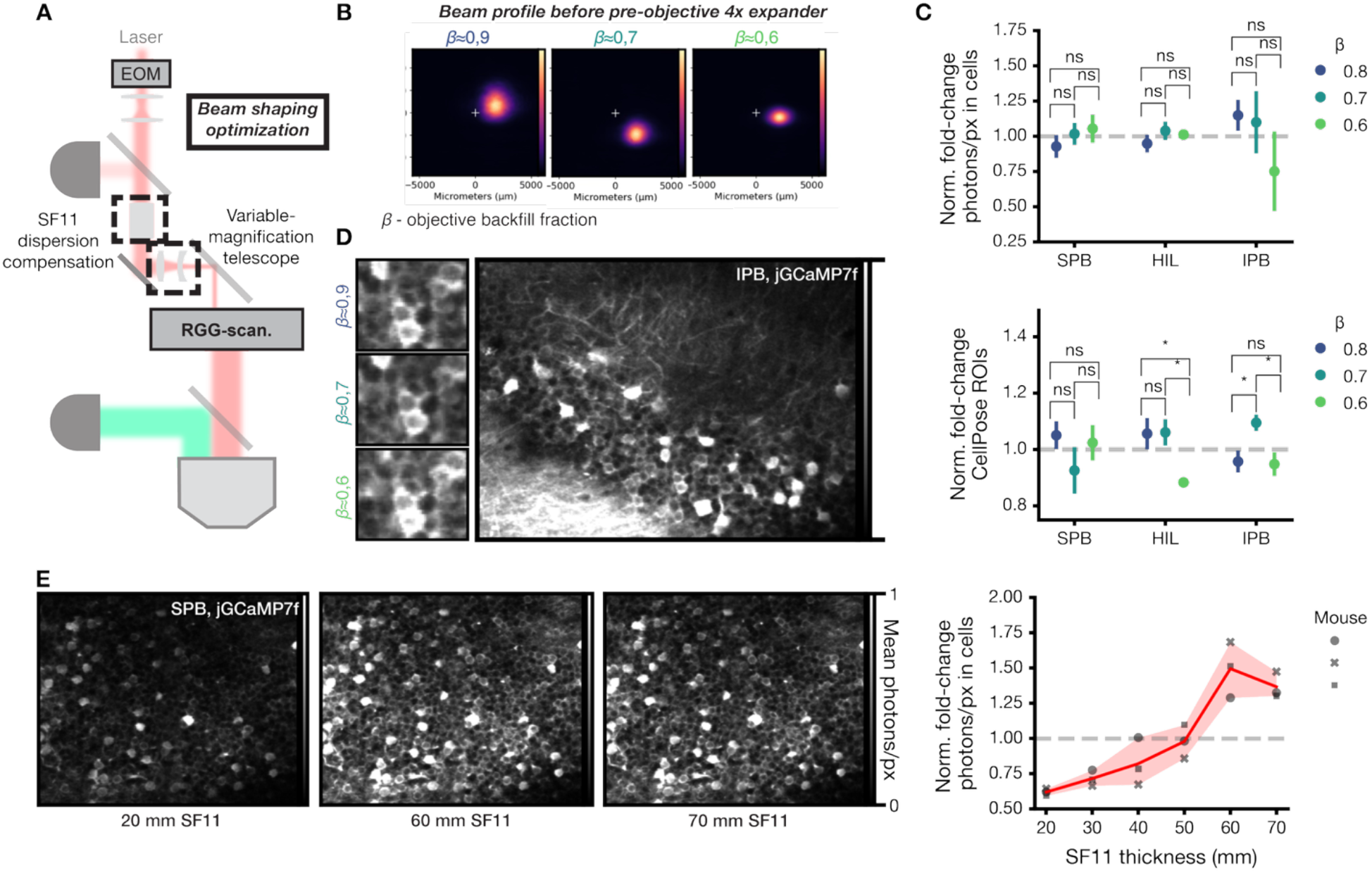
Spatiotemporal excitation pulse optimization. **A)** Overview of introduced optics for spatial pulse shaping (variable-magnification telescope) and temporal pulse shaping (SF11 dispersion compensation). **B)** Beam profiles measured immediately before pre-objective 4x expander with variable magnifications and estimated objective backfilling, β. Acquisition camera was translated between measurements. **C)** Change in number of photons detected per pixel in segmented cells from suprapyramidal blade (SPB), hilus (HIL), and infrapyramidal (IPB) blade across three mice for different objective backfilling values. **D)** Overview of resolution achieved with different objective backfilling proportions. *Left*, example region from the infrapyramidal blade (full region shown for β=0.7). *Full region scale:* 351µm × 260µm. *Right*. Quantification of the number of CellPose-identified ROIs in the different dentate gyrus regions across three mice. **E)** *Left*. Example average images from the SPB of a mouse imaged with variable dispersion *via* introduced SF11 glass thickness. *Scale:* 356µm × 261µm *Right*. Quantification of photons/px in identified cells in the suprapyramidal blades of three mice with variable thicknesses of SF11 glass.

Next, we optimized the temporal compression of our pulse. Glass optics typically introduce positive group-delay dispersion, which stretches ultrashort laser pulses required for efficient 3P illumination. To optimally correct for this, we first applied a strong, over-compensating negative group-delay dispersion to the laser using chirped mirrors at the output of our 1300-nm laser (**Fig. 1A**). Next, downstream in our excitation path, we tested varying thicknesses of positive-chirp SF11 glass while imaging the SPB in three mice. In this way, we empirically optimized our pulse duration (**Fig. 4E**), maximizing the amzount of collected photons received at a given average excitation power.

### Rapid PRED-3P enables functional imaging of dentate granule and mossy cells in the awake and behaving animal

Having optimized both the detection and the shaping of our 3P excitation pulses, we asked whether rapid PRED-3P could be used to successfully image calcium activity during behavior in different regions of the mouse DG. We injected two mice with rAAV-jGCaMP7f throughout the left dorsal DG and implanted a glass cannula over dorsal CA1. Mice were then water-deprived and trained to run for a fixed water reward on a 2-meter cued burlap belt (**Fig. 5A**).^21^ When mice routinely ran more than one lap per minute, the SPB and HIL were imaged for 5 minutes at 20Hz.

**Figure 5.**
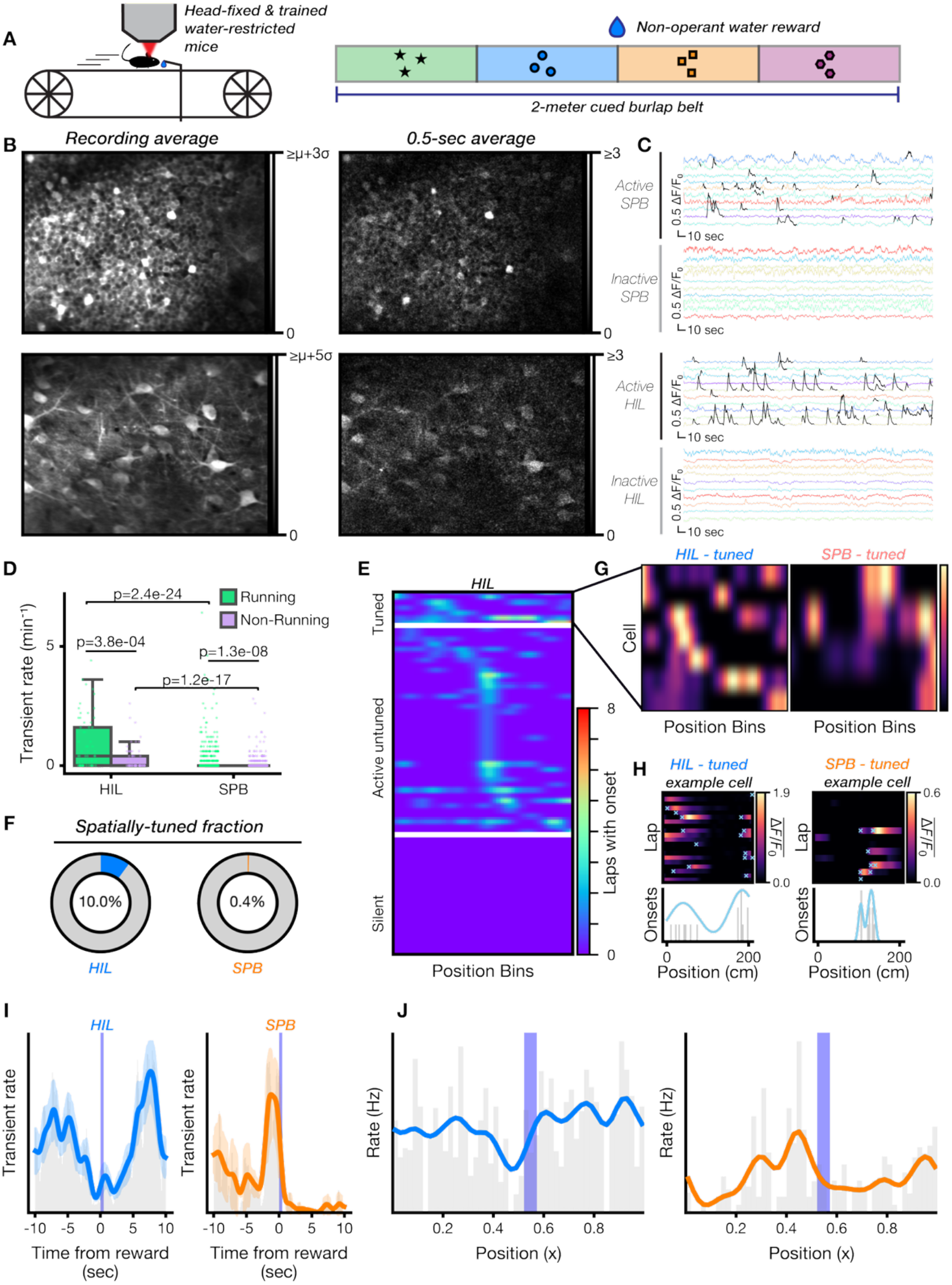
3P functional imaging of calcium activity in the dentate gyrus of awake mice during behavior. **A)** Mice were water restricted, trained to run for water reward on a 2-meter cued burlap belt, and imaged while head fixed during behavior. **B)** *Left*. Example mean FOV from suprapyramidal blade (*SPB*, top) and hilus (*HIL*, bottom) from recording during behavior. Depths: 582-619µm & 620-682µm, respectively. *Right*. Detected signal in photons/pixel of the FOVs to the left during a 0.5-second window. μ= mean pixel value. σ= standard deviation of pixel values. **C)** Example traces from active and inactive cells detected in the SPB and hilus during mouse behavior. Transients are marked in black. **D)** Transient rates during running and non-running epochs in both dentate gyrus regions. **E)** Spatial distribution of transients in hilar mossy cells, separated into spatially tuned, active untuned, and silent groups. *Inset*. **F)** Proportion of SPB granule cells (*left*) and hilar mossy cells (*right*) with statistically significant spatial tuning. **G)** Transient onsets per spatial bin (normalized by maximum bin). **H)** Mean ΔF/F_0_ by spatial bin per lap from identified spatially tuned cells in hilus and SPB. **I)** Peri-stimulus time histograms on reward delivery from the hilus (left) and suprapyramidal blade (right). **J)** Population-level transient rate across spatial bins in HIL and SPB.

A large FOV was captured in every recording, enabling the measurement of approximately 450 putative granule cells (GCs) in SPB and 35 mossy cells (MCs) in HIL per recording (**Fig. 5B, left**). PRED-3P achieves high signal-to-noise (SNR) in our recordings, with >1-photon per pixel obtained per 0.5-second period across cells in the recording (**Fig. 5B, right**).

Calcium transients were successfully measured in both the SPB and the HIL with significantly higher fidelity from the PRED correction (**Fig 5C, Fig. S2**). The measured activity of putative MCs and GCs recapitulates known properties of these cell types.^21,22,27,28^ Firstly, we find that both populations have higher transient rates during running epochs vs. at rest, and that putative MCs in the HIL are substantially more active than GCs, regardless of mobility state (**Fig. 5D**). Next, we find that a much larger proportion of putative MCs is spatially tuned than of GCs (**Fig. 5F**), with both measured proportions agreeing well with previously reported numbers. Most MCs fire at least one transient throughout the recording (**Fig. 5E, left**), and spatially tuned MCs frequently exhibit multiple place fields (**Fig. 5G,H, right**). We find that GCs increase their firing rate in anticipation of reward, whereas putative MCs decrease their firing, agreeing with published results about how these cells behave in response to salient stimuli (**Fig. 5I**). Finally, we find that, as expected, at the population level, putative MCs show no differences in transient rate, except for immediately before the reward zone, whereas GCs interestingly show a ramping like property, with the overall population transient rate increasing with closer proximity to the reward (**Fig. 5J**).

## Discussion

We describe a rapid-repetition-rate fast-scanning three-photon microscope (3PM) implementation that features reduced noise from our single-photon resolution with excitation-fluctuation decoupling (PRED) correction, and improved efficiency from spatiotemporal pulse optimization. Our implementation achieves *bona fide* functional 3PM of neuronal calcium activity in the dorsal dentate gyrus (DG) of awake behaving animals through intact CA1 at frame rates ≥20Hz for the first time.

By tuning our laser repetition rate to 4.14-MHz and combining it with an 8-kHz resonant-galvo-galvo scanner, we achieve 20-28Hz imaging of a ∼250µm × 350µm field-of-view (FOV) containing several hundreds of cells in the cases of the DG supra-(SPB) and infrapyramidal blades (IPB). We demonstrate that 3PM provides superior contrast to 2PM in the DG and enables automated detection of many more cells. We next eliminated most of the excess noise above Poisson shot noise in our detection by using a deeply cooled silicon photomultiplier array (SiPM) and by measuring the excitation pulse powers used to illuminate each pixel in every recorded frame. The SiPM allows conversion of electronic digitizer counts into estimated photon-detection events, reducing electronic noise and in turn, enabling correction of excitation pulse-to-pulse variation using a photon-statistics-based Bayesian correction approach. We further improve our implementation by empirically measuring the optimal objective backfilling proportion and pulse dispersion. Finally, the combination of these improvements allowed us to measure putative granule cells (GCs) in the SPB of the DG and putative mossy cells (MCs) in the hilus (HIL) in mice running for fixed water rewards on a cued burlap belt. Our measurements recapitulate many known functional properties of these cells, even enabling the identification of spatially tuned MCs.

Our 3PM-approach provides a framework to achieve wide-FOV 3PM for the first time at the frame rates required by modern calcium indicators, addressing a major limitation of this imaging method. By implementing our enhancements, 3PM might be used to establish the functional properties of other brain regions during animal behavior, which are otherwise too deep for conventional 2PM, such as layers V & VI of the murine neocortex or non-human primates. Further study of the dorsal DG with this approach is also warranted, as most of the multiphoton imaging in this brain region has been limited to the SPB, with only a few studies or none in the HIL and IPB, respectively. Here, we reliably resolved GCs in the IPB for the first time. Despite the known difference in the physiological and anatomical properties of GCs in the SPB vs. IPB, including their entorhinal inputs,^33,34^ the activity of the infrapyramidal blade has never been imaged during animal behavior due to its depth. Our approach opens a window to do so in future studies.

Even considering the advances that we have made, our excitation approach and detection strategies could be further improved. By implementing adaptive optics,^35-37^ the excitation efficiency of our system could be additionally boosted. Furthermore, the use of specialized objective covers could allow us to collect a larger fraction of the scattered light emitted from the deep brain, thereby enhancing our ability to detect small brightness changes in fluorescent indicators.^38^ Finally, combining 1300-nm excitation with new and emerging infrared genetically encoded calcium indicators might improve signal-to-noise, as emitted photons at these wavelengths should scatter less in biological tissue.^39-40^

Beyond the implications for deep multiphoton imaging, the PRED correction we describe can be utilized in any imaging scenario where small changes in dim photon fluxes need to be detected. High-speed voltage indicator imaging is a prime candidate, as stochastic frame-to-frame fluctuations in laser power could create false-positive spikes and greatly complicate the interpretation of these data. Given that pulse-to-pulse variation has not been routinely reported in these studies, the scope of the problem is unclear, and the PRED approach could be useful for increasing confidence in detected signals.

In summary, our highly optimized rapid PRED-3PM implementation pushes 3P-imaging over the edge for *in vivo* functional imaging of neuronal activity in the neurons too deep for 2P-imaging. By combining our approach with adaptive optics and enhanced collection optics in the future, 3P-imaging would likely become even better for this application. PRED-3P provides a new approach for *in vivo* deep-brain imaging and opens the door for the study of never-before-measured brain regions, including the IPB of the DG.

## Materials and methods

All experiments were conducted in accordance with US National Institutes of Health guidelines and with the approval of the Columbia University Institutional Animal Care and Use Committee. No statistical methods were used to determine sample sizes *a priori*.

### Optical setup

Full description in supplemental discussion #1. Three-photon (3P) excitation light was delivered from a 1300-nm White Dwarf femtosecond laser system (Class V cat. #WD-1300S, Hamburg, Germany). Two-photon (2P) excitation light was delivered from a Chameleon Ultra II Ti:Sapphire laser (Coherent, USA) tuned to 920-nm. 3P- and 2P-beam powers were controlled using lithium tantalate and potassium di-deuterium phosphate electro-optic modulators, respectively (Conoptics, cat.#360-40 and #350-80, USA). A 1% beam pickoff was introduced in the 3P-excitation path, leading to a half-wave plate + polarizing beam splitter, dividing light to a BBO crystal and spectrometer for spectral characterization and to a InGaAs photodiode for pulse-pulse power characterization. Overall negative group-phase dispersion of the 3P beam from the chirped mirrors in the laser was corrected using SF11 glass. The two beams were then combined through a dichroic mirror (2P-beam transmitted, 3P-beam reflected) and steered into a home-built microscope body directing them into a Rapid Multi Region resonant-galvo-galvo scanhead (Vidrio, USA). The beam was then expanded with a 50-mm focal-length scan lens and 200-mm effective focal-length mounted in Plössl configuration and focused using a XLPLN25XWMP2, 25x/1-NA, Olympus objective (Olympus, Japan) dipped in deuterated water. Emitted light was then collected using custom large-aperture 2-inch collection optics and measured using a Thorlabs PMT 2102 or SiPM PDSC21(ThorLabs, USA).

### Beam profile measurement

3P beam profiles after varied expansion were measured using a WinCamD-LCM camera fitted with an 800-nm long-pass filter.

### Mice

For all experiments, we used adult (8-20 weeks) male and female WT mice (C56BL/6). Mice were housed on a 12-h light/12-h dark cycle in groups of 2-5 mice. All procedures were performed under protocols reviewed and approved by Columbia University’s IACUC

### Mouse surgical anesthesia and analgesia

Anesthesia was induced using 4% isoflurane for 4-minutes and maintained at 1.5% isoflurane in 95% oxygen during the duration of the procedure. Body temperature was maintained using a heating pad during the procedure and during the post-operative recovery periods. After induction, mice were head-fixed using a stereotactic instrument (Kopf Instruments, USA). Subcutaneous meloxicam and local subcutaneous bupivacaine were given for analgesia. Post-operatively, 1ml of saline was delivered subcutaneously.

### Stereotactic adeno-associated virus (AAV) injections

After induction of anesthesia and delivery of analgesia, a midsagittal incision of the scalp was performed. Adhesive tissue to the bone was aspirated and the bregma and lambda skull sutures were identified. The head positioned was flattened using the stereotactic instrument such that the axial positions of bregma and lambda were ≤100µm. Burr holes were made using a dental drill at 1,8mm caudal, 1,1mm lateral + 2,2mm caudal, 1,5mm lateral from bregma. 50nl of AAV9-hSyn-jGCaMP8s-WPRE (Addgene, #162374-AAV9) or AAV1-CamKII-jGCaMP7f (Hen lab, Columbia University) were delivered at each site at depths of 2mm, 1,8mm, 1,6mm, and 1,2mm from the surface of the brain. The scalp was re-sealed using buried simple interrupted stitches with 5-0 nylon suture.

### Cannula implantation

Pre-operatively, 3-mm #1,5 coverslips were attached to 3-mm diameter imaging cannulae using UV-curable adhesive. Cannulae were sterilized in 70% EtOH. Cannulae implants were performed at least three days after AAV injection. After head fixation, the scalp overlying the parietal and the posterior half of the frontal bones was removed. Residual adhesive tissue on the bone was aspirated, and the skull was scored with a razor blade. A 3-mm punch biopsy was used to perform a craniotomy containing both burr holes. Dura and cortex were aspirated slowly with continuous irrigation of ice-cold phosphate buffered saline until coronal corpus callosum fibers were visible under magnification. An imaging cannula was then lowered into the craniotomy and positioned flat using the stereotactic instrument. Cannulae were secured first by Vetbond and subsequently with C&B Metabond (Parkell). A custom titanium headpost was then secured at the juncture of the parietal and interparietal bones with Metabond and a layer of dental acrylic.

### Behavior training

At least 7 days after cannula implantation, animals were acclimated to head fixation on a burlap belt over two days with 20 minutes of head fixation on each. Animals were then water-restricted and given 20 minutes on a burlap belt with a lickport delivering 20 randomly located water rewards per 2-meter lap. As the animals learned to run for water, the reward frequency was reduced to 10, 5, 3, 2, and finally 1 reward per lap until animals ran at least one lap per minute. At this point, they were switched to a cued burlap belt with a single reward delivery per lap at a fixed position (1-meter mark). Recordings were done once animals ran at least one lap per minute on the cued belt.

### Imaging acquisition details

All recordings were taken through deuterated water. For the 2P/3P comparison, the same FOVs were imaged with both modes for two minutes. For 2P acquisition (since the laser features a higher repetition rate), the whole field was acquired at 56 Hz, rather than the two serially acquired tiles at 28 Hz for 3P. 2P recordings were taken before 3P measurements. All 3P recordings were average power-matched to their 2P counterparts. For behavioral imaging, each field of view was imaged at 20Hz and was recorded for 5 minutes.

### Automated cell detection

CellPose-SAM^41^ was used to automatically segment cells from the recording average of motion-corrected recordings. Segmented regions of interest (ROIs) were accepted as putative granule cells if their area was ≥40% that expected of cells with a diameter of at least 10,5µm in the suprapyramidal or infrapyramidal blades of the dentate gyrus, and as putative mossy cells with a diameter of at least 22µm in the dentate hilus. For 2PM vs. 3PM comparison (Fig. 2), CA1 PNs were identifying by counting masks with areas ≥50-pixels and ≤250 pixels. IPB and SPB GCs were identified ≥50-pixels and ≤180-pixels. HIL MCs were identified ≥180 pixels.

### Recording pre-processing

Stochastic resonant scanner resampling correction (supplemental discussion #2) was first applied to each recorded tile and its corresponding laser pulse power recording. Rigid motion correction was performed with Suite2P^42^ and then repeated using the average of the motion-corrected recording as the initial seed. Rigid registration shifts were applied to the laser channel. PRED correction was applied to each rigid motion-corrected tile (see supplemental discussion #3). Nonrigid motion correction was then performed on each tile separately. Rigid and non-rigid motion shifts were then used to reconstruct expanded motion-corrected tiles (including regions occupied in only a few frames). Overlapping regions on expanded left and right tiles were identified and aligned. A potential brightness gradient across the seam from imperfect beam centering on the x-galvanometer was corrected by rescaling the brightness of the left and right tiles to the mean using a constant scaling factor. Cells were segmented as above. Traces were extracted and neuropil was decontaminated by modeling traces as true signal, contamination from neighboring cells, and contamination from neuropil. The latter two components were removed. Baselines were estimated on the raw and corrected traces using the CWT-BR algorithm.

### Transient detection

Indicator transients were identified from baseline-subtracted neuropil-corrected traces using the constrained FOOPSI implementation in the CaImAn analysis package.^43^ Peaks with an amplitude under 2-standard deviations above the mean or <0.3-seconds were rejected. Baselines were re-fit excluding peaks via a running median on remaining frames. Peak identification was then repeated iteratively three times (with standard deviation cutoffs of 2, 2.5, and 3 in that order). Composite events (composed of multiple transients) were separated if in the FOOPSI-denoised traces they featured peak prominences >0.25 and the smaller peak was >0.1 the height of the larger peak. Transient candidates were analyzed against the noise by sampling 200 non-transient segments and computing a z-score and p-value of the power-mean of the true events vs. the null distribution. Events with p>0.01 were considered “false,” re-labeled as background, and the process was iterated until convergence. Final events with z-score > 6 were kept for downstream analysis. Traces were smoothed using a Savitzky-Golay filter for visualization.

### Spatial tuning calculation

Categorization of cells as spatially tuned was performed similarly to previously reported.^24^ Briefly, Skaggs spatial information content was computed from transient onset times using 2, 4, 5, 8, 10, 20, 25, and 100 spatial bins. We then performed 100,000 random circular shuffles of the transient onset times and re-computed the information content and subtracted the mean of this distribution from the measured spatial information contents. We thus chose a single information content estimate from the recorded transient onsets and from the shuffles as that which is maximal across all bin sizes tested for each. The spatial tuning p-value was taken as the fraction of shuffles in which the spatial information content exceeded that of the true onsets. Cells with p-values < 0.05 were accepted as spatially-tuned.

## Author contributions

AL and TG conceived the original project. AL and DSP designed the study and supervised the project. TSM, AL, and DSP wrote the manuscript. TSM collected data. TSM, DSP, and AL analyzed the data. AN built the initial microscope. TSM, DSP, EK, and TG modified it and made subsequent improvements. TSM and DSP implemented the PRED correction. TSM and EK performed animal surgeries, and behavioral training.

