## supplementary information for "Photon-Resolved Excitation-Denoised (PRED) Three-Photon Imaging Improves Detection of Neuronal Activity in Awake and Behaving Mice"

### *Supporting Information*

#### Supplemental figures

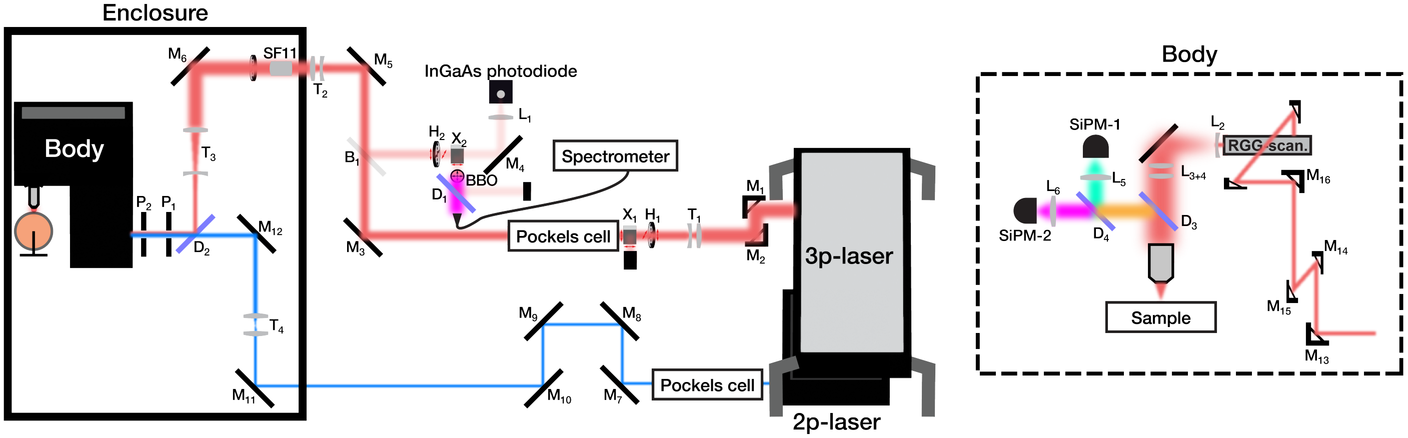


***B*** - beamsampler ***D*** - dichroic ***H*** - half-wave plate ***L*** - lens ***M*** - mirror ***P*** - pinhole ***S*** - shutter ***T*** - telescope ***X*** - polarizing beam-splitter

**Fig. S1** ***Optical diagram of the combined 2P-/3P-imaging setup.*** The red beam illustrates the 3P-beam path, and the blue shows the 2P-path.

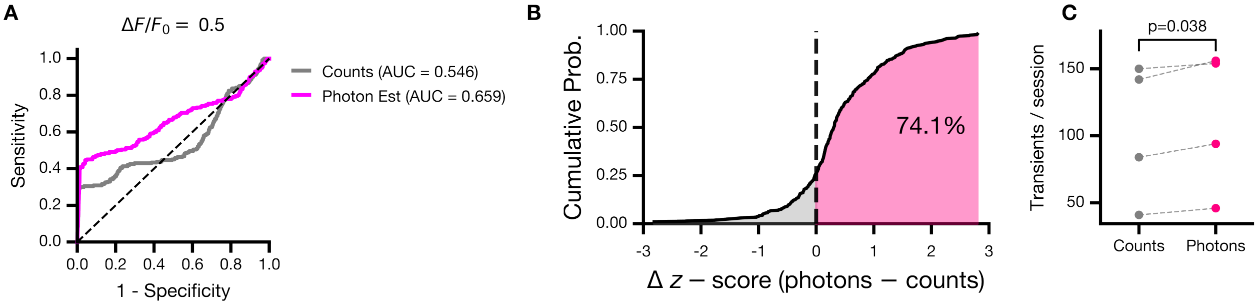


**Fig. S2** ***PRED correction vs. transient detection.*** **A)** Receiver-operating-curves for PRED-corrected vs uncorrected recording of a uniform slide with artificial GCaMP transients injected into the recording. **B)** Cumulative distribution function plot of jGCaMP7f transients from the dentate hilus and suprapyramidal blade identified during animal behavior/locomotion. Transients identified in both the PRED-corrected and uncorrected data were extracted and their z-score is compared between the two channels. **C)** Comparison of transients identified per session in PRED-corrected and uncorrected *in vivo* recordings. *p*-value from paired t-test.

#### Supplemental discussions

##### §1 - Scanning correction.

Given the sinusoidal velocity of a resonant scanner in time, the scanner position for a bidirectional "back-and-forth" is given by:

$$x_{\mathrm{scan}}[t] = \frac{1}{2}\Big(1 - \mathbf{cos}\big[\frac{\pi}{\tau}t\big]\Big) \tag{1}$$

where:

$$t = \text{time}$$
$$\tau = \text{unidirectional line period}$$
$$x_{\text{scan}} : \; [0, 1]$$

In an acquisition, $t_0$ is set by a synchronization pulse from the resonant scanner that marks the beginning of a new bidirectional line period. A phase offset may exist between this pulse and the true beginning of the line period. This phase offset may be corrected during acquisition, yet a non-zero amount may remain, modifying equation (1) to:

$$x_{\mathrm{scan}}[t] = \frac{1}{2}\Big(1 - \mathbf{cos}\big[\frac{\pi}{\tau}(t + \theta)\big]\Big)$$

where:

$$\theta = \text{scanner-sync clock to line-period start delay}$$

Assuming that laser pulses occur at a regular interval, the scanner position may be re-formulated as a function of laser pulse number:

$$x_{\mathrm{scan}}[i] = \frac{1}{2}\Big(1 - \mathbf{cos}\Big[\frac{\pi}{\tau}\big(\frac{i}{r} + \theta\big)\Big]\Big)$$

where:

$$i = \text{laser pulse number}$$
$$r = \text{laser repetition rate}$$

This can be further generalized to account for a non-zero delay between the starting scanner synchronization clock pulse and the succeeding laser pulse time:

$$x_{\mathrm{scan}}[i] = \frac{1}{2}\Big(1 - \mathbf{cos}\Big[\frac{\pi}{\tau}\big(\frac{i}{r} + \theta + \psi_f\big)\Big]\Big)$$

where:

$$\psi_f = \text{initial scanner-sync clock to laser-pulse delay}$$

*NB* - $\psi_f$ is a function of the frame number, $f$, as any frame rate which is an imperfect multiple of the laser repetition rate will cause a frame-to-frame jitter in this phase.
Finally the equation can be transformed from scanner space to physical space:

$$x[i] = \frac{w\rho}{2f_{\mathsf{x}}}\Big(1 - \mathbf{cos}\Big[\frac{\pi}{\tau}\big(\frac{i}{r} + \theta + \psi_f\big)\Big]\Big) - \frac{w}{2}\Big(\frac{1}{f_{\mathsf{x}}} - 1\Big)$$

or further simplified:

$$x[i] = \frac{w\rho}{2f_{\mathsf{x}}}\Big(f_{\mathsf{x}} - \cos\Big[\frac{\pi}{\tau}\big(\frac{i}{r} + \theta + \psi_f\big)\Big]\Big) \tag{2}$$

$$w = \text{width of the imaged region in pixels}$$
$$\rho = \text{pixel width}$$
$$f_{\mathsf{x}} = \text{spatial-fill fraction}$$
$$\tau = \text{unidirectional line period}$$
$$i = \text{laser pulse number}$$
$$r = \text{laser repetition rate}$$
$$\theta = \text{scanner-sync clock to line-period start delay}$$
$$\psi_f = \text{initial scanner-sync clock to laser-pulse delay for frame } f \text{ (\textbf{??})}$$

All the values in equation (2) are known parameters of the imaging system, outside of $\theta$ and $\psi$, which must be estimated. To determine the true spatial position recorded in each pixel, the position is first calculated for every laser pulse within the recording (including those that fall outside the fill fraction). Next the scanner synchronization clock times are identified as:

$$t_{\text{sync}} = 2n\tau$$

where:

$$n = 0, 1, 2, ...$$

and laser pulse times are identified as:

$$t_{\text{pulse}} = \frac{i}{r} + \psi_0$$

where:

$$i = 0, 1, 2, ...$$

$$\psi_0 = \text{scanner-sync clock to line-period start delay at recording start}$$

Then, the first laser pulse of each line is identified by the following relation:

$$\tau(l + \frac{1 - f_{\text{t}}}{2}) \leq t_{\text{pulse}}$$

where:

$$l = \text{line number: } 0, 1, 2, ...$$

and the next $w$ pulses are kept, where $w$ is the number of pixels per line. Now that the indices of the laser pulses, $i$, that appear in the image have been solved and together with equation (2), $\theta$ and $\psi_f$ can be estimated from well-aligned imaging data as follows.

The frame start time is defined as the nearest resonant scanner cycle to the nominal frame time:

$$t_f = 2\tau \cdot \text{round}\left(\frac{f/f_s}{2\tau}\right)$$

The laser phase at frame $f$ is then given by evaluating the laser clock at $t_f$:

$$\begin{aligned} \psi_f &= (rt_f + \psi_0) \bmod 1 \\ &= \left(\psi_0 + 2\tau r \cdot \text{round}\left(\frac{f/f_s}{2\tau}\right)\right) \bmod 1 \end{aligned} \tag{3}$$

where $\psi_0$ is the initial laser-scanner phase offset at the start of acquisition.

$\psi_0$ and $\theta$ are finally fit using the following algorithm:

**Algorithm: Scanning phase optimizations**

1. A subset of up to 30 frames from the most highly-correlated 3-seconds of recorded frames are extracted and averaged in time.
2. Bounds-constrained L-BFGS-B minimization of the loss metric is performed starting from the grid search optimum.
3. A localized $7 \times 7$ grid polish is conducted around the L-BFGS-B solution, followed by a final L-BFGS-B refinement step to secure the optimized parameters.

$$\text{Loss} = -\text{Coherence} + 0.1 \times \left(\frac{\text{EO}_{L1}}{\text{stdev}(R)}\right) \tag{4}$$

$$\text{Coherence} = \frac{1}{M} \sum_{k \in \text{ROI}} \left| \frac{1}{N} \sum_{n=1}^{N} \frac{\mathcal{F}[R]_{n,k}}{|\mathcal{F}[R]_{n,k}| + \epsilon} \right| \tag{5}$$

$$\text{EO}_{L1} = \frac{1}{H} \sum_{m=1}^{H} |R_{2m} - R_{2m+1}| \tag{6}$$

#### §2 - Photon-resolution excitation-denoising (PRED) correction.

##### §2.1 - Template fitting

Initial templates are fit using the following algorithm:

**Algorithm: Initial template fitting**

1. A pixel histogram is generated from 1-million random pixels below the 99th-percentile.
2. The histogram is smoothed with a Gaussian kernel ($\sigma$=0.5).
3. Pixel values below the center position are reflected across the $x$ position of the peak.
4. Peaks are identified in SciPy ($\mathrm{find\_peaks}$)
5. A linear-regression is fit to calculated initial peak centers.
6. Gaussian functions are positioned on the regression line at every integer peak number and are fit for amplitude and universal (shared) $\sigma$.

Next, the 0-photon pedestal and the 1-photon template are fit as below:

**Algorithm: 0- and 1-photon template fitting**

1. The pixel histogram is cut off at the center position of the initial 0-photon peak.
2. Pixel values below the center position are reflected across the $x$ position of the peak.
3. A Gaussian function is fit for amplitude and $\sigma$ to this zero-photon pedestal.
4. A Gaussian with the same $\sigma$ as above is placed at the peak position for the initial 1-photon template and fit for amplitude.
5. The 1-photon template is extracted as the original Gaussian at values above the peak position and as the smoothed residual between the histogram and the 0- and 1-photon Gaussians to the left of the peak.

After this, higher-number photon template shapes are generated, as follows:

$$T_n[i] = \begin{cases} \sum_m T_{n-1}[m]\; T_1[i-m], & i \leq i_{\text{peak}} \\ T_n[i_{\text{peak}}] \cdot \exp\left(-\frac{1}{2}\left(\frac{i-i_{\text{peak}}}{\sigma}\right)^2\right), & i > i_{\text{peak}} \end{cases} \tag{7}$$

$$i = \frac{c - c_0}{\Delta_i} \tag{8}$$

where:

$$\begin{aligned} n &= \text{photon number: } 2, 3, 4, \ldots \\ \sigma &= \text{standard deviation from 0-photon pedestal} \\ \mathrm{c} &= \text{analogue-to-digital converter counts} \\ \Delta_i &= \text{ADC count bin width} \end{aligned}$$

Finally, each template is normalized to have AUC = 1:

$$\bar{T}_n[i] = T_n[i] / \sum_i T_n[i] \tag{9}$$

##### §2.2 - Dark-statistics fitting

***Dark count modeling***

When the excitation laser power is zero, any detected events come from one of two sources:

1. *Dark counts*. Random avalanche diode discharges.
2. *Crosstalk*. Avalanche diode discharges triggered by other discharges.

We first calculate weights $w'_n$ to $\bar{T}_n$ for $n = 0, 1, 2$ to best fit the pixel histogram, $h[i]$, in the dark region.

$$\{w_n\} = \mathbf{argmin}_{w_n \forall n \leq 2} \Big[ \sum_{n=0}^{2} w'_n \cdot \bar{T}_n[i] - h[i] \Big] \tag{10}$$

We assume that events in the inter-ROI region of the image where the electro-optic modulator is "zeroed" are overwhelmingly dark counts or crosstalk. Given that dark counts are random & independent events that occur at a constant rate, the dark count probability follows a Poisson distribution, where:

$$P_{\text{dc}}(n) = \frac{\lambda_{\text{dc}}^n \mathrm{e}^{-\lambda_{\text{dc}}}}{n!}$$

Since crosstalk events must be, by definition, triggered by another discharge event, 1-photon events must be dark counts. Thus:

$$\frac{P_{\text{dc}}[1]}{P_{\text{dc}}[0]} = \frac{\sum_{\mathrm{i}}(w'_1 \cdot \bar{T}_1[i])}{\sum_i(w'_0 \cdot \bar{T}_0[i])} = \frac{\lambda_{\text{dc}} \mathrm{e}^{-\lambda_{\text{dc}}}}{\mathrm{e}^{-\lambda_{\text{dc}}}}$$

Simplifying via the normalization of the templates and rearranging:

$$\lambda_{\text{dc}} = \frac{w'_1}{w'_0}$$

In summary, the probability and rate of dark count events are:

$$\boxed{\begin{aligned} P_{\text{dc}}(n) &= \frac{\lambda_{\text{dc}}^n \mathrm{e}^{-\lambda_{\text{dc}}}}{n!} \\ \lambda_{\text{dc}} &= \frac{w'_1}{w'_0} \\ &\text{where } w_n \text{ via (10)} \end{aligned}} \tag{11}$$

***Crosstalk modeling***

Multi-photon events in the dark region may either be:

1. $n$ independent dark counts
2. Some number of dark counts $n - i$ with $i$ crosstalk events

Unlike dark counts, crosstalk events are neither independent, nor random (depending on preceding events), and thus do not follow a simple Poisson distribution. Instead, crosstalk can be effectively modeled as a *branching* Poisson process, where each avalanche diode discharge has a small probability of producing "offspring" crosstalk discharge(s), which in turn can produce their own discharges, and so on. The number of total discharges given a single "parent" discharge follows a Borel distribution:

$$P_{\text{ct + parent}}[n] = \mathbf{Borel}[n|\lambda_{\text{ct}}] = \frac{(\lambda_{\text{ct}} n)^{n-1} \mathrm{e}^{-\mathrm{n}\lambda_{\text{ct}}}}{n!}$$

This can be generalized from a single parent discharge to the Borel-Tanner distribution, given $r$ total parent discharges:

$$P_{\text{ct + parent}}[n] = \mathbf{Borel-Tanner}[n|r, \lambda_{\text{ct}}] = \frac{r}{n} \cdot \frac{(n\lambda_{\text{ct}})^{n-r}}{(n-r)!} e^{-n\lambda_{\text{ct}}}$$

Given $r$ parents, the probability of $n$ crosstalk events can be solved

$$P_{\text{ct}}[n|r,\lambda_{ct}] = P_{\text{ct+parent}}[n+r] = \frac{r}{n+r} \cdot \frac{\left((n+r)\lambda_{\text{ct}}\right)^n}{n!} \mathrm{e}^{-(n+r)\lambda_{\text{ct}}} \tag{12}$$

Now, $\lambda_{\text{ct}}$ must be solved. Let $P_k$ be the probability of one discharge producing $k$ offspring discharges. The expected number of crosstalk discharges per single discharge is therefore:

$$\lambda_{\text{ct}} = \sum_{k=0}^{\infty} kP_k$$

The expected number of first-generation crosstalk discharges (i.e., those from the initial avalanche diode discharges "generations") is:

$$d_1 = \lambda_{\text{ct}} d_0$$

Each first-generation crosstalk discharge may produce its own second generation discharges, which can produce third generation discharges, etc.

$$\begin{aligned} d_0 &= d_0 \\ d_1 &= \lambda_{\text{ct}} \cdot d_0 = \lambda_{\text{ct}} d_0 \\ d_2 &= \lambda_{\text{ct}} \cdot d_1 = \lambda_{\text{ct}}^2 d_0 \\ &\vdots \\ d_g &= \lambda_{\text{ct}}^g d_0 \end{aligned}$$

Crosstalk discharges are defined as those produced after generation $0$, so:

$$d_{\text{ct}} = d_{\text{total}} - d_0 = \sum_{g=1}^{\infty} d_g = d_0 \Big(\sum_{g=0}^{\infty} \lambda_{\text{ct}}^g - 1\Big)$$

Assuming that each discharge produces <1 crosstalk discharge on average, the geometric series in the first term converges, giving:

$$d_{ct} = d_0 (\frac{1}{1-\lambda_{\text{ct}}} - 1)$$

or:

$$d_{ct} = d_0 (\frac{\lambda_{\text{ct}}}{1-\lambda_{\text{ct}}})$$

Given that $\lambda_{\text{ct}}$ is the expected number of discharges from a single avalanche diode discharge, this can be re-stated as the rate of multiphoton discharges in excess of that expected from the dark-count rate per single-photon discharge in the dark region, i.e.:

$$\lambda_{\text{ct}} = \frac{\sum_{d=2}^{\infty} (P[d] - P_{\text{dc}}[d])}{P[1]}$$

or in terms of the normalized templates:

$$\lambda_{\text{ct}} = \frac{\sum_{d=2}^{\infty} \left(w'_d \cdot \sum_i (\bar{T}_d[i]) - \frac{\lambda_{\text{dc}}^d \mathrm{e}^{-\lambda_{\text{dc}}}}{d!}\right)}{\lambda_{\text{dc}} \mathrm{e}^{-\lambda_{\text{dc}}}} = \frac{\sum_{d=2}^{\infty} \left(w'_d - \frac{\lambda_{\text{dc}}^d \mathrm{e}^{-\lambda_{\text{dc}}}}{d!}\right)}{\lambda_{\text{dc}} \mathrm{e}^{-\lambda_{\text{dc}}}}$$

In summary, the probability distribution and rate of crosstalk events, given $r$ parent events are:

$$\boxed{\begin{aligned} P_{\text{ct}}[n|r] &= \frac{r}{n+r} \cdot \frac{\left((n+r)\lambda_{\text{ct}}\right)^n}{n!} \mathrm{e}^{-(n+r)\lambda_{\text{ct}}} \\ \lambda_{\text{ct}} &= \frac{\sum_{d=2}^{\infty} \left(w'_d - \frac{\lambda_{\text{dc}}^d \mathrm{e}^{-\lambda_{\text{dc}}}}{d!}\right)}{\lambda_{\text{dc}} \mathrm{e}^{-\lambda_{\text{dc}}}} \end{aligned}}$$

*NB*- In practice, since it is exceedingly rare for multi-photon crosstalk events in the dark region, in order to avoid over-influence from electronic noise, the crosstalk rate is approximated as:

$$\lambda_{\text{ct}} \approx \frac{w'_2 - \frac{\lambda_{\text{dc}}^2 \mathrm{e}^{-\lambda_{\text{dc}}}}{2}}{\lambda_{\text{dc}} \mathrm{e}^{-\lambda_{\text{dc}}}} \tag{13}$$

##### §2.3 - Generating detector-corrected templates matrix

After fitting the dark count rate $\lambda_{\text{dc}}$ and crosstalk parameter $\lambda_{\text{ct}}$ from the dark region, we aim to estimate the overall distribution of emitted photon counts per pixel in the recording.

Given $j$ initial events (emissions or dark counts), the probability of observing $n$ total events after crosstalk amplification is given by the Borel-Tanner distribution via (2). The kernel for the overall contribution of dark counts and resultant crosstalk to the distribution is then obtained by convolving the dark count probability distribution (11) with the crosstalk distribution:

$$K_{\text{combined}}[n] = \sum_{j=0}^{n} \frac{\lambda_{\text{dc}}^{j} e^{-\lambda_{\text{dc}}}}{j!} \cdot \frac{j}{n} \cdot \frac{(n\lambda_{\text{ct}})^{n-j}}{(n-j)!} e^{-n\lambda_{\text{ct}}}$$

The combined kernel is used to construct a Toeplitz transformation matrix, which functionally encodes how an ideal photon count distribution would be distorted by the properties of the detector (dark counts and crosstalk):

$$\mathbb{M}_{\text{trans}} = \textbf{Toeplitz}[K_{\text{combined}}]$$

The final template matrix is obtained by applying the transformation to the ideal templates:

$$\mathbb{T} = \mathbb{M}_{\text{trans}} \times \bar{\mathbb{T}}$$

$$\text{where } \bar{\mathbb{T}}[n,i] = \bar{T}_n[i]$$

Now, to estimate the overall distribution of emitted photon counts for any given pixel throughout the recording, the distortion-corrected templates must be linearly combined to best fit the histogram of ADC values over all frames for the pixel, $h_i \equiv \vec{h}$. To solve this system of linear equations, the Moore-Penrose pseudo-inverse of the template matrix is computed as:

$$\mathbb{T}^{+} = (\mathbb{T}^{\text{T}}\mathbb{T})^{-1}\mathbb{T}^{\text{T}}$$

The histogram is modeled as a linear combination of templates:

$$\vec{h} \approx \mathbb{T} \cdot \vec{w}$$

The least-squares solution for photon count weights is thus:

$$\vec{w} = \mathbb{T}^{+} \cdot \vec{h} \tag{14}$$

##### §2.4 - Pixel-wise baseline brightness estimation

For a pixel in a 3-photon excitation process, the intrinsic brightness $b$ is defined as the photon flux per laser power cubed:

$$\lambda = b \cdot L^3 \tag{15}$$

where:

$\lambda =$ photon flux
$b =$ brightness
$L =$ excitation laser power

Given that the laser power and brightness can change as a function of time, at each frame $t$ with laser power $L_t$, the expected photon count distribution is Poisson:

$$P(n|b_t, L_t) = \frac{(b_t L_t^3)^n e^{-b_t L_t^3}}{n!} \tag{16}$$

If calcium transients are rare, we can assume that the brightness is almost always at its baseline, $\bar{b}$ (and thus a constant). Thus, the overall photon count probability distribution for a set of $T$ frames is given by:

$$\bar{P}(n|\bar{b}, \vec{L}_t) = \frac{1}{T} \sum_{t=0}^{T} \frac{(\bar{b} L_t^3)^n e^{-\bar{b} L_t^3}}{n!}$$

Given that the photon weights $\vec{w}$ represent the probability distribution of different emitted photon numbers for a given pixel, a likelihood function is defined in terms of brightness, using (14) and (2):

$$\mathcal{L}(\bar{b}) = \prod_n \left( \frac{1}{T} \sum_{t=0}^{T} \bar{P}(n|\bar{b}, \vec{L}_t) \right)^{w_n}$$

Baseline brightness, $\bar{b}$, for a given pixel is solved by maximizing the log-likelihood function, giving:

$$\bar{b} = \mathbf{argmax}_{\bar{b}} \sum_n \left[ w_n \mathbf{log} \left[ \frac{1}{T} \sum_{t=0}^{T} \bar{P}(n|\bar{b}, \vec{L}_t) \right] \right] \tag{17}$$

In biological samples imaged *in vivo*, motion during the recording violates the assumption that a given pixel contains the same emitter(s) in every frame. When applying the correction to *in vivo* recordings, a preliminary rigid motion correction is first performed, and the top 25% of frames most-correlated with the average of the motion corrected recording are used for estimating this parameter.

**§2.5 - Photons per pixel probability versus time.**

We first ask, what is the probability of detecting $\gamma_t$ photons at time $t$, given $c_t$ ADC counts?

By Bayes' theorem:

$$P[\gamma_t|c_t] = P[c_t|\gamma_t] \cdot \frac{P[\gamma_t]}{P[c_t]} \tag{18}$$

**Lemma I: Probability of detecting $c_t$ ADC counts given $\gamma$ emitted photons**

Goal: solve the following term:

$$P[c|\gamma]$$

We marginalize the conditional probability over $d$ detector discharges. Given that $\gamma$ discharges come from emitted photons, $d \geq \gamma$:

$$P[c|\gamma] = \sum_{d \geq \gamma} P[c|d] \cdot p[d|\gamma]$$

The first term is the probability density of getting $c$ counts from $d$ discharges, i.e. $\bar{T}_d[c]$. The second term can be re-phrased as the probability of having $k \equiv d - \gamma$ discharges from dark counts or crosstalk.

$$P[c|\gamma] = \sum_{d \geq \gamma} \bar{T}_d[c] \sum_{i=0}^{k} P_{\text{dc}}[i|\lambda_{\text{dc}}] \cdot P_{\text{ct}}[(k-i)|(\gamma+i), \lambda_{\text{ct}}]$$

Substituting in the terms from (11) and (2), gives:

$$P[c|\gamma] = \sum_{d \geq \gamma} \bar{T}_d[c] \sum_{i=0}^{k} \left( \frac{\lambda_{\text{dc}}^i e^{-\lambda_{\text{dc}}}}{i!} \right) \cdot \left( \frac{\gamma+i}{d} \cdot \frac{\left(d\lambda_{\text{ct}}\right)^{(k-i)}}{(k-i)!} e^{-(\gamma+i)\lambda_{\text{ct}}} \right)$$

Given that $\binom{k}{i} = \frac{k!}{i!(k-i)!}$:

$$P[c|\gamma] = \sum_{d \geq \gamma} \bar{T}_d[c] \frac{\mathrm{e}^{-(\lambda_{\text{dc}}+\gamma\lambda_{\text{ct}})}}{d \cdot k!} \sum_{i=0}^{k} \binom{k}{i} \lambda_{\text{dc}}^i (d\lambda_{\text{ct}})^{k-i} (\gamma+i) \mathrm{e}^{-i\lambda_{\text{ct}}}$$

Define $\Lambda \equiv \lambda_{\text{dc}} \mathrm{e}^{-\lambda_{\text{ct}}}$ and substitute

$$P[c|\gamma] = \sum_{d \geq \gamma} \bar{T}_d[c] \frac{\mathrm{e}^{-(\lambda_{\text{dc}}+\gamma\lambda_{\text{ct}})}}{d \cdot k!} \left( \gamma \sum_{i=0}^{k} \binom{k}{i} \Lambda^i (d\lambda_{\text{ct}})^{k-i} + \sum_{i=0}^{k} i \binom{k}{i} \Lambda^i (d\lambda_{\text{ct}})^{k-i} \right)$$

Using the identity for the binomial expansion $(x+y)^k = \sum_{i=0}^{k} \binom{k}{i} x^i y^{k-i}$ and recognizing $x \frac{\mathrm{d}}{\mathrm{d}x}(x+y)^k$ :

$$P[c|\gamma] = \sum_{d \geq \gamma} \bar{T}_d[c] \frac{\mathrm{e}^{-(\lambda_{\mathrm{dc}} + \gamma \lambda_{\mathrm{ct}})}}{d \cdot k!} \left( \gamma (\Lambda + d\lambda_{\mathrm{ct}})^k + \Lambda k (\Lambda + d\lambda_{\mathrm{ct}})^{k-1} \right) \tag{19}$$

Thus, altogether we have:

$$P[c|\gamma] = \sum_{d \geq \gamma} w_d \bar{T}_d[c] \tag{20}$$

where :

$$w_d = \frac{\mathrm{e}^{-(\lambda_{\mathrm{dc}} + \gamma \lambda_{\mathrm{ct}})}}{d \cdot k!} \big( \Lambda + d\lambda_{\mathrm{ct}} \big)^{k-1} \cdot \Big( \gamma (\Lambda + d\lambda_{\mathrm{ct}}) + k\Lambda \Big)$$

$$\Lambda \equiv \lambda_{\mathrm{dc}} \mathrm{e}^{-\lambda_{\mathrm{ct}}}$$

$$k \equiv d - \gamma$$

The probability of detecting $\gamma_t$ photons may be marginalized over brightness:

$$P[\gamma_t] = \int_{b=0}^{\infty} \mathrm{d}b \cdot \underbrace{P[\gamma_t|b]}_{\mathbf{Poisson}[\gamma_t; bL_t^3]} \cdot p[b]$$

The probability of detecting $c_t$ digitizer counts can be marginalized over $\gamma_t$ discharges coming from emitted photons at time $t$:

$$P[c_t] = \sum_{\gamma_t} P[c_t|\gamma_t] \cdot P[\gamma_t]$$

Marginalizing the second term over brightness gives:

$$P[c_t] = \sum_{\gamma_t} P[c_t|\gamma_t] \int_{b=0}^{\infty} \mathrm{d}b \cdot \underbrace{P[\gamma_t|b]}_{\mathbf{Poisson}[\gamma_t; bL_t^3]} \cdot p[b]$$

Thus, the probability is:

$$P[\gamma_t|c_t] = \frac{P[c_t|\gamma_t] \cdot \int_{b=0}^{\infty} \mathrm{d}b \cdot \mathbf{Poisson}[\gamma; bL_t^3] \cdot p[b]}{\sum_{\gamma} P[c_t|\gamma] \cdot \int_{b=0}^{\infty} \mathrm{d}b \cdot \mathbf{Poisson}[\gamma; bL_t^3] \cdot p[b]} \tag{21}$$

$P[c_t|\gamma_t]$ via equation (20)

**Case 1:** $p[b] = \boldsymbol{\delta}[b - \bar{b}]$

Via the sifting property of the Dirac delta function, (21) reduces to

$$P[\gamma_t|c_t] = \frac{P[c_t|\gamma_t] \cdot \mathbf{Poisson}[\gamma; \bar{b}L_t^3]}{\sum_{\gamma} P[c_t|\gamma] \cdot \mathbf{Poisson}[\gamma; \bar{b}L_t^3]} \tag{22}$$

$P[c_t|\gamma_t]$ via equation (20)

**Case 2:** *Brightness $b$ is almost always fixed at its baseline $\bar{b}$ (at rest) but can occasionally increase (during activity).*

We model the probability density of brightness as a mixture model between a delta function at its baseline with an exponential tail:

$$p[b] = (1-\epsilon)\boldsymbol{\delta}[b - \bar{b}] + \frac{\epsilon}{\beta} \mathrm{e}^{-\frac{b-\bar{b}}{\beta}} \text{ for b} \geq \bar{\text{b}}$$

The integral becomes:

$$\int_{b=0}^{\infty} \mathrm{d}b \cdot \mathbf{Poisson}[\gamma; bL_t^3] \cdot p[b] = (1-\epsilon) \int_{b=0}^{\infty} \mathrm{d}b \cdot \mathbf{Poisson}[\gamma; bL_t^3] \cdot \boldsymbol{\delta}[b-\bar{b}] + \frac{\epsilon}{\beta} \int_{b=\bar{b}}^{\infty} \mathrm{d}b \cdot \mathbf{Poisson}[\gamma; bL_t^3] \cdot \mathrm{e}^{-\frac{b-\bar{b}}{\beta}}$$

$$= (1-\epsilon) \cdot \mathbf{Poisson}[\gamma; \bar{b}L_t^3] + \frac{\epsilon}{\beta} \int_{b=\bar{b}}^{\infty} \mathrm{d}b \frac{(bL_t^3)^\gamma}{\gamma!} \mathrm{e}^{-(bL_t^3 + \frac{b-\bar{b}}{\beta})}$$

The second term can be further reduced:

$$\frac{\epsilon}{\beta} \int_{b=\bar{b}}^{\infty} \mathrm{d}b \frac{(bL_t^3)^\gamma}{\gamma!} \mathrm{e}^{-(bL_t^3 + \frac{b-\bar{b}}{\beta})} = \frac{\epsilon}{\beta} \int_{b=\bar{b}}^{\infty} \mathrm{d}b \frac{(bL_t^3)^\gamma}{\gamma!} \mathrm{e}^{-b(L_t^3 + \frac{1}{\beta}) + \frac{\bar{b}}{\beta}}$$

$$= \epsilon \frac{L_t^{3\gamma}}{\beta\gamma!} \mathrm{e}^{\frac{\bar{b}}{\beta}} \int_{b=\bar{b}}^{\infty} \mathrm{d}b \cdot b^\gamma \cdot \mathrm{e}^{-kb}$$

$$\text{where : } k \equiv L_t^3 + \frac{1}{\beta}$$

Recognizing the integral as the upper-incomplete Gamma function, the second term reduces to:

$$\frac{\epsilon}{\beta} \int_{b=\bar{b}}^{\infty} \mathrm{d}b \frac{(bL_t^3)^\gamma}{\gamma!} \mathrm{e}^{-(bL_t^3 + \frac{b-\bar{b}}{\beta})} = \epsilon \frac{L_t^{3\gamma} \mathrm{e}^{\frac{\bar{b}}{\beta}}}{\beta\gamma! \cdot k^{\gamma+1}} \mathbf{\Gamma}[\gamma+1, k\bar{b}]$$

Thus, altogether the probability becomes

$$P[\gamma_t|c_t] = \frac{P[c_t|\gamma_t] \cdot \big((1-\epsilon) \cdot \mathbf{Poisson}[\gamma_t; \bar{b}L_t^3] + \epsilon \cdot \xi[\gamma_t]\big)}{\sum \gamma P[c_t|\gamma] \cdot \big((1-\epsilon) \cdot \mathbf{Poisson}[\gamma; \bar{b}L_t^3] + \epsilon \cdot \xi[\gamma]\big)} \tag{23}$$

*where* :

$\epsilon \approx$ likelihood for having brightness above baseline

$$\xi[\gamma] = \frac{L_t^{3\gamma} \mathrm{e}^{\frac{\bar{b}}{\beta}}}{\beta\gamma! \cdot k^{\gamma+1}} \mathbf{\Gamma}[\gamma+1, k\bar{b}]$$

$\beta \approx$ scale of the supra-baseline deviations relative to baseline

$k \equiv L_t^3 + \beta^{-1}$

$P[c_t|\gamma_t]$ $via$ equation (20)

**§2.6 - Expected underlying brightness vs. time.**

The number of emitted photons will vary with the excitation laser power. The brightness of the fluorescent indicator, however, is invariant to this and only changes with the actual underlying biological phenomenon being sensed (e.g., a calcium transient). Thus, we aim to calculate the expected brightness of each pixel at a given time, $\hat{b}_t$.

Given the expected photon number $\hat{\gamma}_t$ at time $t$, by definition of the expected value:

$$\bar{\gamma}_t = \sum_{\gamma_t=0}^{\infty} \gamma_t \cdot P[\gamma_t] = \sum_{\gamma_t=0}^{\infty} \gamma_t \cdot \mathbf{Poisson}[\gamma_t | b_t L_t^3]$$

Evaluating the expected value yields:

$$\hat{\gamma}_t = \sum_{\gamma_t=1}^{\infty} \gamma_t \cdot \frac{(b_t L_t^3)^{\gamma_t}}{\gamma_t!} \mathrm{e}^{-b_t L_t^3}$$

$$= (b_t L_t^3) \mathrm{e}^{-b_t L_t^3} \sum_{\gamma_t=1}^{\infty} \frac{(b_t L_t^3)^{\gamma_t - 1}}{(\gamma_t - 1)!}$$

$$= (b_t L_t^3) \mathrm{e}^{-b_t L_t^3} \sum_{k=0}^{\infty} \frac{(b_t L_t^3)^k}{k!}$$

Recognizing the sum as the Maclaurin series for the exponential function:

$$\hat{\gamma}_t = (b_t L_t^3)\mathrm{e}^{-b_t L_t^3} \cdot \mathrm{e}^{b_t L_t^3} = b_t L_t^3$$

Therefore the expected brightness at time $t$ is:

$$\hat{b}_t = \hat{\gamma}_t \frac{1}{L_t^3}$$

In our measurement, we have more information about the probability of emitted photon number $\gamma_t$ from the number of ADC counts, $c_t$. Thus, substituting the expected photon number gives the final answer:

$$\boxed{\hat{b}_t = \frac{1}{L_t^3} \sum_{\gamma} \gamma \cdot P[\gamma | c_t]} \tag{24}$$

where $P[\gamma | c_t]$ is calculated via (23).
